# Delineating the rules for structural adaptation of membrane-associated proteins to evolutionary changes in membrane lipidome

**DOI:** 10.1101/762146

**Authors:** Maria Makarova, Maria Peter, Gabor Balogh, Attila Glatz, James I. MacRae, Nestor Lopez Mora, Paula Booth, Eugene Makeyev, Laszlo Vigh, Snezhana Oliferenko

## Abstract

Membrane function is fundamental to life. Each species explores membrane lipid diversity within a genetically predefined range of possibilities. How membrane lipid composition in turn defines the functional space available for evolution of membrane-centered processes remains largely unknown. We address this fundamental question using related fission yeasts *Schizosaccharomyces pombe* and *Schizosaccharomyces japonicus*. We show that unlike *S. pombe* that generates membranes where both glycerophospholipid acyl tails are predominantly 16-18 carbons long, *S. japonicus* synthesizes unusual ‘asymmetrical’ glycerophospholipids where the tails differ in length by 6-8 carbons. This results in stiffer bilayers with distinct lipid packing properties. Retroengineered *S. pombe* synthesizing the *S. japonicus*-type phospholipids exhibits unfolded protein response and downregulates secretion. Importantly, our protein sequence comparisons and domain swap experiments indicate that transmembrane helices co-evolve with membranes, suggesting that, on the evolutionary scale, changes in membrane lipid composition may necessitate extensive adaptation of the membrane-associated proteome.

## Results and Discussion

The fission yeasts *Schizosaccharomyces pombe* and *Schizosaccharomyces japonicus* show remarkable differences in fundamental membrane-centered processes such as establishment of polarity and mitotic nuclear envelope (NE) remodeling [1, 2]. The different strategies of managing the NE have been linked to the distinct regulation of lipid synthesis during the cell cycle in the two species [3, 4]. To analyze membrane lipid compositions of the two sister species, we performed shotgun ESI-MS/MS analysis of total cellular lipid extracts (Supplemental Table 1). The most abundant membrane lipids were GPLs, defined by the polar headgroups and two fatty acyl chains (FA) at the *sn-1* and *sn-2* positions of the glycerol backbone (Fig. 1A). We observed subtle differences in the abundance of major GPL classes, including phosphatidylcholine (PC), phosphatidylethanolamine (PE), phosphatidylinositol (PI) and phosphatidylserine (PS) (Fig. 1B). We also detected some variation in the abundance of the minor GPL classes and sphingolipids (Supplemental Fig. 1A and Supplemental Table 1). *S. japonicus* contained higher amounts of storage triacylglycerols, whereas the cellular levels of sterol esters were comparable between the two species (Supplemental Table 1).

**Fig. 1.**
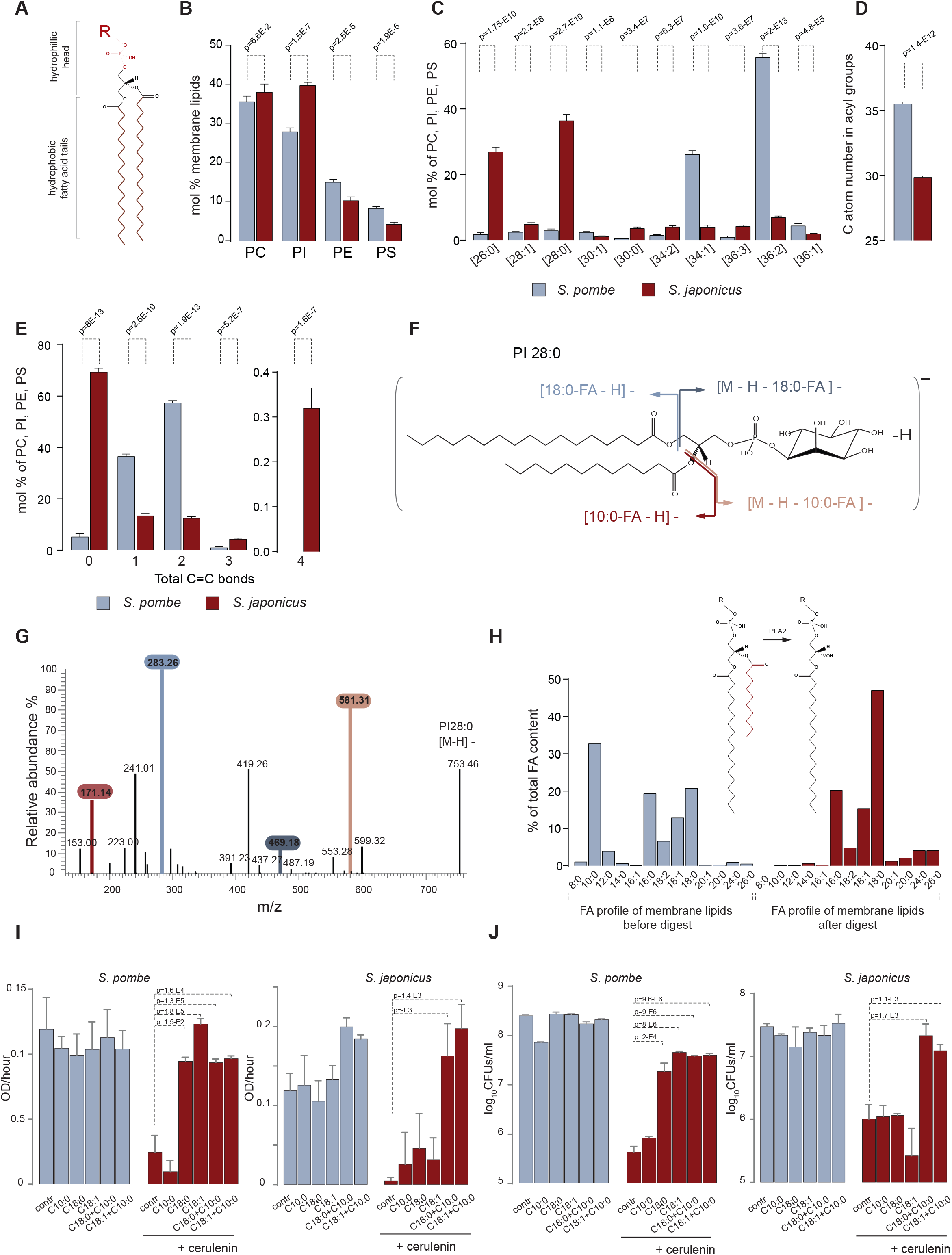
*S. japonicus* relies on production of membrane glycerophospholipids exhibiting pronounced acyl chain asymmetry. (**A**) Generic structure of a GPL molecule containing two hydrophobic FA chains and a hydrophilic head group. (**B**) Relative abundance of the four main GPL classes (PC, PI, PE, and PS) in *S. pombe* and *S. japonicus*. (**C**) Average FA composition of the PC, PI, PE, and PS GPLs in the two species. The categories shown are defined based on the total number of carbon atoms : total number of double bonds in acyls. (**D**) Average combined FA length in PC, PI, PE and PS the two species. (**E**) Comparison of FA saturation in PC, PI, PE and PS the two species. The GPLs are categorized into four groups depending on the total number of the double bonds in the two acyls. (**F**) A diagram of the 28:0 PI species abundant in *S. japonicus*. Colors and arrows indicate ions formed upon fragmentation. (**G**) A typical mass spectrum of the C28:0 PI species from the *S. japonicus* lipid extract indicating ions generated upon fragmentation. Colors match the indicated fragments in panel (**G**). (**H**) Mass spectrometry analysis of FA content in GPLs before and after PLA2 treatment, which removes FAs specifically from the *sn*-2 position. Note that C10:0 is not recovered after digestion indicating that it is located at the *sn*-2 position. (**I**) Comparison of growth rates of *S. pombe* and *S. japonicus* cultures under indicated conditions. Growth rates were calculated for an exponential region of the growth curve as a change in OD_595_ per hour. **(J)** Survival of *S. pombe* and *S. japonicus* cells grown in EMM supplemented with indicated FAs loaded on BSA. Colony forming units (CFUs) per ml were counted after 48 hours of growth. (**B**, **C**, **D**, **E**) Shown are the mean values ± SD (*n* = 5). *p* values derived from the unpaired parametric t-test. (**I-J**) Shown are the mean values ± SD (*n* = 3). *p* values derived from the unpaired parametric t-test.

Remarkably, we observed differences in the FA chain composition between *S. pombe* and *S. japonicus*. In line with previous reports, the most abundant molecular species in major GPL classes (PC, PE, PI and PS) in *S. pombe* were 36:2 and 34:1, where the first number indicates the combined length of FA tails and the second, total number of double bonds in FA tails (Fig. 1C, Supplemental Tables 2-3 and [5]). The size profile of *S. pombe* GPLs was thus similar to the lipidome of the model budding yeast *Saccharomyces cerevisiae* [6]. However, *S. japonicus* GPLs exhibited a markedly distinct chemical composition, with abundant 26:0 and 28:0 molecular species (Fig. 1C-D and Supplemental Tables 2-3). Such a trend towards lower molecular weight species was present in every analyzed lipid class (Supplemental Table 2-3).

In addition to differences in total length, phospholipid acyl chains were more saturated in *S. japonicus* compared to *S. pombe* (Fig. 1E and Supplemental Table 3). Despite this, we identified some polyunsaturated GPLs in *S. japonicus*, which were not detected in *S. pombe*, suggesting differences in desaturase activity (Fig. 1E). This is consistent with the *S. japonicus* genome encoding a delta-12 desaturase, in addition to the delta-9 desaturase Ole1 [7], common to both fission yeasts and other eukaryotic groups [8].

In order to elucidate the structure of the lower molecular weight GPL species, we performed fragmentation analysis of the major classes of GPLs in *S. japonicus* (Fig. 1F and Supplemental Tables 4-5). The resulting mass spectra showed that C16 and C18 long chain fatty acyls were frequently found in combination with the medium-chain fatty acyl C10:0, forming asymmetrical structures, with C10:0 located in the *sn-2* position of the glycerol backbone (see data for an abundant PI species C28:0 in Fig. 1F-G, with full interpretation of data in Supplemental Table 4; data for the other major GPLs are presented in Supplemental Table 5). We confirmed that C10:0 was found invariably in the *sn-2* position by analyzing the products of the *sn-2* specific phospholipase A2 (PLA2) digestion (Fig. 1H). The proportion of such highly asymmetrical structures, where the two chains differed in length by 6-8 carbon atoms, varied between phospholipid classes, reaching ∼90% for PI (Supplemental Fig. 1B and Supplemental Table 5). We concluded that *S. japonicus* synthesizes a large proportion of highly asymmetrical membrane GPLs.

The abundance of asymmetrical C10:0-containing GPLs in *S. japonicus* suggested that this medium chain FA might be essential for *S. japonicus* physiology. To test this directly, we inhibited endogenous FA synthesis with cerulenin [9] and supplemented both *S. pombe* and *S. japonicus* with exogenous FAs of different lengths and saturation. As expected, cell growth and survival of both species decreased drastically upon cerulenin treatment. The addition of exogenous long-chain fatty acids (LCFAs) such as oleic (C18:1) or stearic acid (C18:0) was sufficient to rescue both parameters in *S. pombe* (Fig. 1I-J and Supplemental Fig. 1C). In the same experimental setup, LCFAs failed to rescue cerulenin-induced defects in growth and survival of *S. japonicus* cells. C10:0 alone was also insufficient, but when it was added to cerulenin-treated *S. japonicus* cultures in combination with either C18:1 or C18:0, the growth and survival defects were largely rescued (Fig. 1I-J and Supplemental Fig. 1D). Together, these results demonstrate that *S. japonicus* physiology relies on the presence of both C10:0 and long chain fatty acids.

Differences in lipid structure may bear on biological functions by altering membrane’s properties. One of the fundamental biophysical parameters is the bending rigidity, which reflects how much energy is needed to deform the membrane from its intrinsic curvature. This energy reflects the bilayer stiffness [10]. We have generated giant unilamellar vesicles (GUVs), which are thought to mimic the biophysical properties of the cellular membranes, from purified *S. pombe* or *S. japonicus* GPLs. These GUVs were then subjected to osmotic stress to induce membrane fluctuations (Fig. 2A). Analysis of thermal membrane fluctuations demonstrated that the bending rigidity parameter of membranes assembled from the *S. japonicus* phospholipid mixture was approximately two-fold higher than that originating from *S. pombe* GUVs (Fig. 2A). This indicated that GPL asymmetry and/or increased FA saturation in *S. japonicus* might contribute to higher bilayer stiffness.

**Fig. 2.**
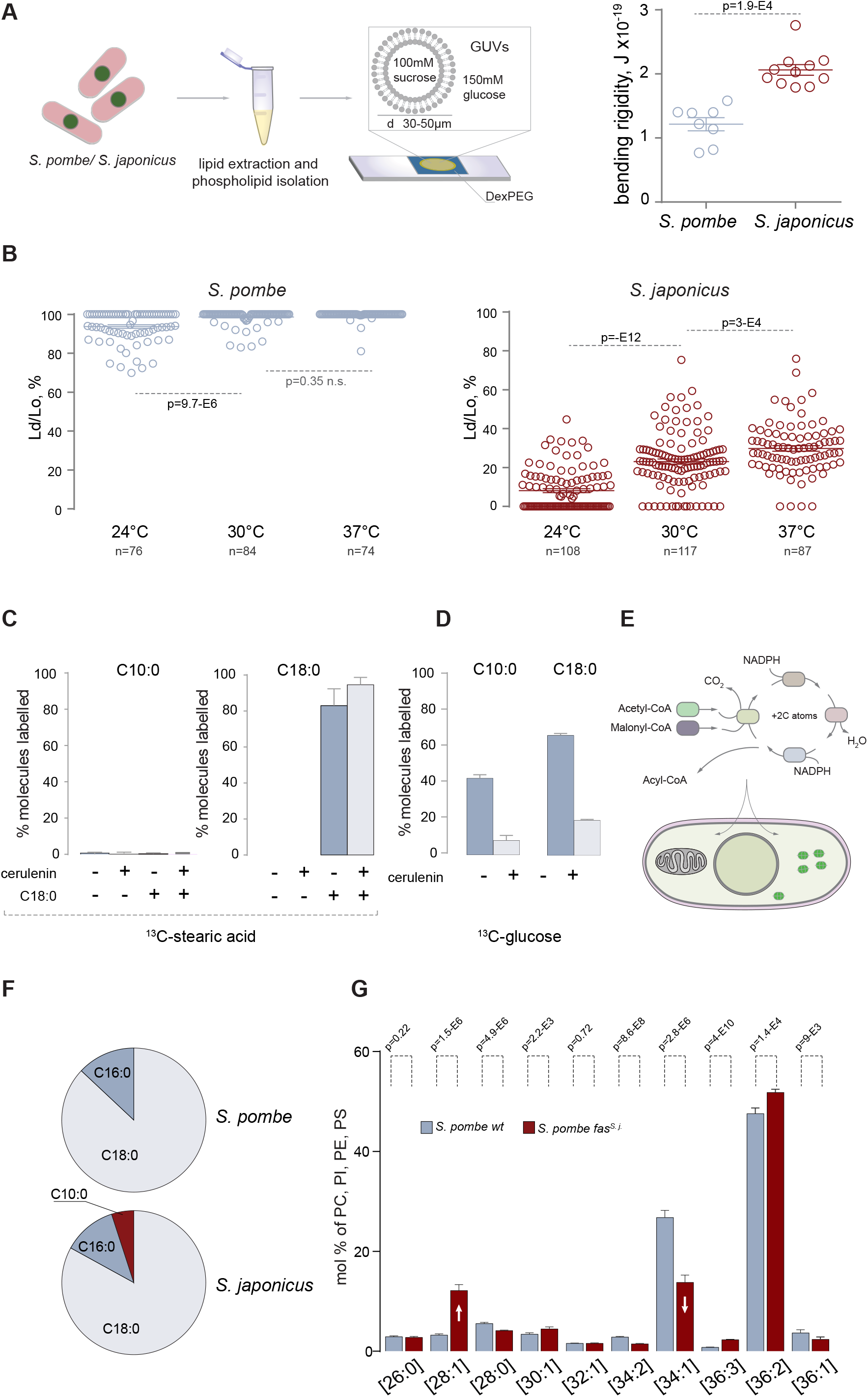
C10:0 is synthesized in cytosol by the FAS complex and contributes to the higher membrane stiffness and lipid packing. (**A**) *Left*, in a typical experiment GUVs in a 30-50 μm range were obtained by hydrogel formation and thermal fluctuations were analyzed by light microscopy. Bending rigidities were calculated using fluctuation analysis. *Right*, One-dimensional scatter plot summarizing the results of these experiments. *n* = 8 and 11 for *S. pombe* and *S. japonicus*, respectively. (**B**) One-dimensional scatter plots summarizing the results of the temperature-dependent Lo and Ld phase separation in GUVs made of *S. pombe* (*left*) and *S. japonicus* (*right*) phospholipids. The Ld/Lo ratios were calculated from the measurements of the areas of distinct intensities at the middle plane of GUVs. Numbers of GUV measurements are indicated underneath each column. (**A**-**B**) *In vitro* experiments were carried out with GPL fractions isolated from the total lipid extracts. Means ± SD are indicated and the *p* values are derived from the Kolmogorov-Smirnov test. (**C**) A plot summarizing the GC-MS analysis of ^13^C-labeled C10:0 and C18:0 FAs extracted from *S. japonicus* cells grown in the presence of the U-^13^C-C18:0 for four hours. 10 μM cerulenin was added when indicated. Shown are the means of percentages of ^13^C-labeled FAs ± SD (n = 6). (**D**) A plot describing the results of the GC-MS analysis of ^13^C-labeled C10:0 and C18:0 FAs extracted from *S. japonicus* cells grown in the presence of the U-^13^C-glucose for four hours. 10 μM cerulenin was added when indicated. Shown are the means of percentages of ^13^C-labeled FAs ± SD (n = 6). (**E**) A schematic of the FA synthesis in eukaryotes. FAs are synthesized from the priming substrate acetyl-CoA and the elongating substrate malonyl-CoA. Each cycle is comprised of two reduction and one dehydration steps resulting in acyl chain growth by two carbon atoms. FA synthesis reactions in eukaryotes can be executed by two pathways, localized either in the cytosol or mitochondria. (**F**) A pie chart summarizing the results of shotgun mass spectrometry analysis of fatty acyl-CoA products generated *in vitro* by purified cytosolic FA synthases from *S. pombe* and *S. japonicus*. Shown are percentages of acyl-CoA species identified in each reaction. (**G**) A graph representing the average molecular species profiles for the four indicated lipid classes in *S. pombe* wild type and *S. pombe fas^s. j.^* cells. Phospholipids are specified by the total carbon atoms: total double bonds in acyls. Note an increase in the 28:1 at the expense of 34:1 species. Shown are the mean values ± SD (*n* = 4 for the wild type and 5 for *fas^s. j.^* cells). *p* values derived from the unpaired parametric t-test.

Lipid packing is another important biophysical parameter that determines membrane properties [11]. We therefore examined the lipid lateral segregation in the bilayer of *S. pombe* and *S. japonicus* GUVs by examining the fluorescence intensity of the lipid tracer Fast DiO, which has a strong preference for the liquid disordered (Ld) over the liquid ordered (Lo) phase [12]. Our analyses revealed that GUVs of *S. pombe* at 24°C were predominantly in the Ld phase with some GUVs showing coexistence of Ld and Lo phases (Fig. 2B and Supplemental Fig. 2A). The proportion of the Lo phase decreased with raise in temperature, with most *S. pombe* membranes becoming fully disordered at 37°C. In contrast, GUVs assembled from *S. japonicus* glycerophospholipids exhibited pronounced membrane order at 24°C. Increasing temperature resulted in the growth of Ld domains in *S. japonicus*-derived membranes but phase coexistence was maintained at 37°C, which is at the higher end of the physiologically relevant temperature range (Fig. 2B and Supplemental Fig. 2A). Taken together, these results demonstrate that the divergence in glycerophospholipid structures between the two fission yeasts may have a profound effect on the biophysical properties of their membranes, resulting in higher membrane stiffness and a distinct phase behavior in *S. japonicus*.

We then set out to understand biosynthetic origins of the C10:0 abundance in *S. japonicus*. Using metabolic labeling followed by gas chromatography-mass spectrometry (GC-MS), we ruled out the possible shortening of LCFAs as a source of this medium-chain fatty acid (MCFA). Inhibition of endogenous FA synthesis by cerulenin and subsequent supplementation with stably labeled stearic acid (^13^C-C18:0) did not result in the detection of ^13^C-labeled C10:0 (Fig. 2C), which was in line with our observation that supplementation with LCFAs was not sufficient to rescue *S. japonicus* growth upon fatty acid synthase (FAS) inhibition (Fig. 1I,J). This suggested that C10:0 likely originated through *de novo* FA synthesis. Indeed, analysis *of S. japonicus* cells grown in the presence of ^13^C-labeled glucose revealed that the rates of synthesis were comparable between C10:0 and other FAs and that all FA production was inhibited by the FAS inhibitor cerulenin (Fig. 2D and Supplemental Fig. 2B).

Fungi and animals encode two pathways for *de novo* fatty acid biosynthesis – located in the mitochondria or the cytosol (Fig. 2E). In mitochondria, several enzymes execute each step of FA synthesis separately [13]. Deletion of three genes (*htd2Δ*, *mct1Δ* and *etr1Δ*), whose products are predicted to execute essential steps of mitochondrial FA synthesis in *S. japonicus*, did not result in changes to the relative abundance of C10:0, suggesting that the mitochondrial pathway was not involved in C10:0 production in this organism (Supplemental Fig. 2C).

The cytosolic fatty acid synthase is a multi-enzyme complex containing all catalytic activities required for the reaction sequence as discrete functional domains [14]). In most fungi, it is comprised of two large polypeptide chains. To address the role of cytosolic FAS in C10:0 production, we purified FAS complexes from either *S. pombe* or *S. japonicus* and reconstituted FA synthesis *in vitro*. The FAS activity was monitored by measuring the malonyl-CoA- and acetyl-CoA-dependent rates of NADPH oxidation (Supplemental Fig. 2D and [15]). Mass spectrometry analyses of the reaction products revealed that, in contrast to the *S. pombe* FAS that produced only C18:0 and some C16:0, the purified *S. japonicus* enzyme was able to synthesize C10:0 alongside the LCFAs, albeit with a lower yield compared to the steady-state abundance of C10:0 *in vivo* (Fig. 2F and Supplemental Table 6).

We wondered if the genetic exchange of the *S. pombe* cytosolic FAS to its *S. japonicus* counterpart would be sufficient to alter the spectrum of FA products and, as a result, structural composition of glycerophospholipids. The FAS complex is composed of α and β subunits forming an α_6_β_6_ oligomer [14]. We substituted the open reading frames (ORFs) encoding both the α and the β subunits (*fas2* and *fas1*, respectively) in *S. pombe* with their *S. japonicus* orthologs, maintaining the native regulatory contexts unchanged. Remarkably, we observed a pronounced increase in lower molecular weight species (mostly C28:1) for all major GPL classes in the engineered *S. pombe* strain (Fig. 2G, Supplemental Fig. 2E-H and Supplemental Table 7). Fragmentation analyses showed that such lipids contained C18 FAs in the *sn-1* and C10:0 in the *sn-2* positions of the glycerol backbone (as an example, see MS/MS spectra for PI C28:1 in Supplemental Fig. 2I). Interestingly, an increase in the C28:1 species coincided with decreased abundance of C34:1 lipids containing C18:1 and C16:0 chains (Fig. 2G). This suggested that the *S. japonicus* FAS was sufficient to increase C10:0 content at the expense of C16:0, and that C10:0 was incorporated in place of a saturated long FA chain into *S. pombe* glycerophospholipids. Additionally, we detected a reduced GPL to protein ratio (Supplemental Table 7) and subtle differences in membrane lipid composition in *S. pombe* cells expressing the transplanted *S. japonicus* FAS, including a slightly increased PC/PE ratio (Supplemental Fig. 2J-K).

Although *S. japonicus* FAS was sufficient to synthesize C10:0 both *in vitro* and *in vivo*, the relative amounts of C10:0 were lower compared to *S. japonicus* cells. This suggests that other factors may enhance the efficiency of C10:0 synthesis in that organism. The acyl-CoA-binding protein (ACBP) was shown to be involved in FA termination and transport and, when in excess, to shift the FAS product spectrum towards MCFAs *in vitro* [16]. However, neither deletion nor replacement of the gene encoding the *S. japonicus* ACBP (*acb1*) with its *S. pombe* ortholog led to significant changes in the relative amount of C10:0 as compared to the wild type *S. japonicus* (Supplemental Fig. 4L). This indicated that the generation of C10:0 in *S. japonicus* does not rely on ACBP activity and other factors will need to be examined in future studies.

Our data so far indicated that the cytosolic FA synthase in *S. japonicus* have diverged from its *S. pombe* counterpart to produce a bimodal distribution of products peaking at C10:0 and C18:0. Interestingly, replacement of the *S. pombe* FAS with the corresponding FAS complex from *S. japonicus* (*fas^s. j.^*) increased population doubling times and decreased cell survival across the physiological range of temperatures. These phenomena were particularly pronounced at 24°C in both rich and minimal media (Fig. 3A-B and Supplemental Fig. 3A). *S. pombe fas^s. j.^* cultures grew at comparable rates in glucose-rich and respiratory media (Supplemental Fig. 3B) and consumed oxygen similarly to the wild type (Supplemental Fig. 3C), indicating that mitochondrial function was not overtly affected by FAS substitution. Addition of exogenous oleic acid (C18:1) to *S. pombe fas^s. j.^* cells rescued cellular growth rates, which suggested that the enhanced C10:0 production could be at root of slow growth (Supplemental Fig. 3D).

**Fig. 3.**
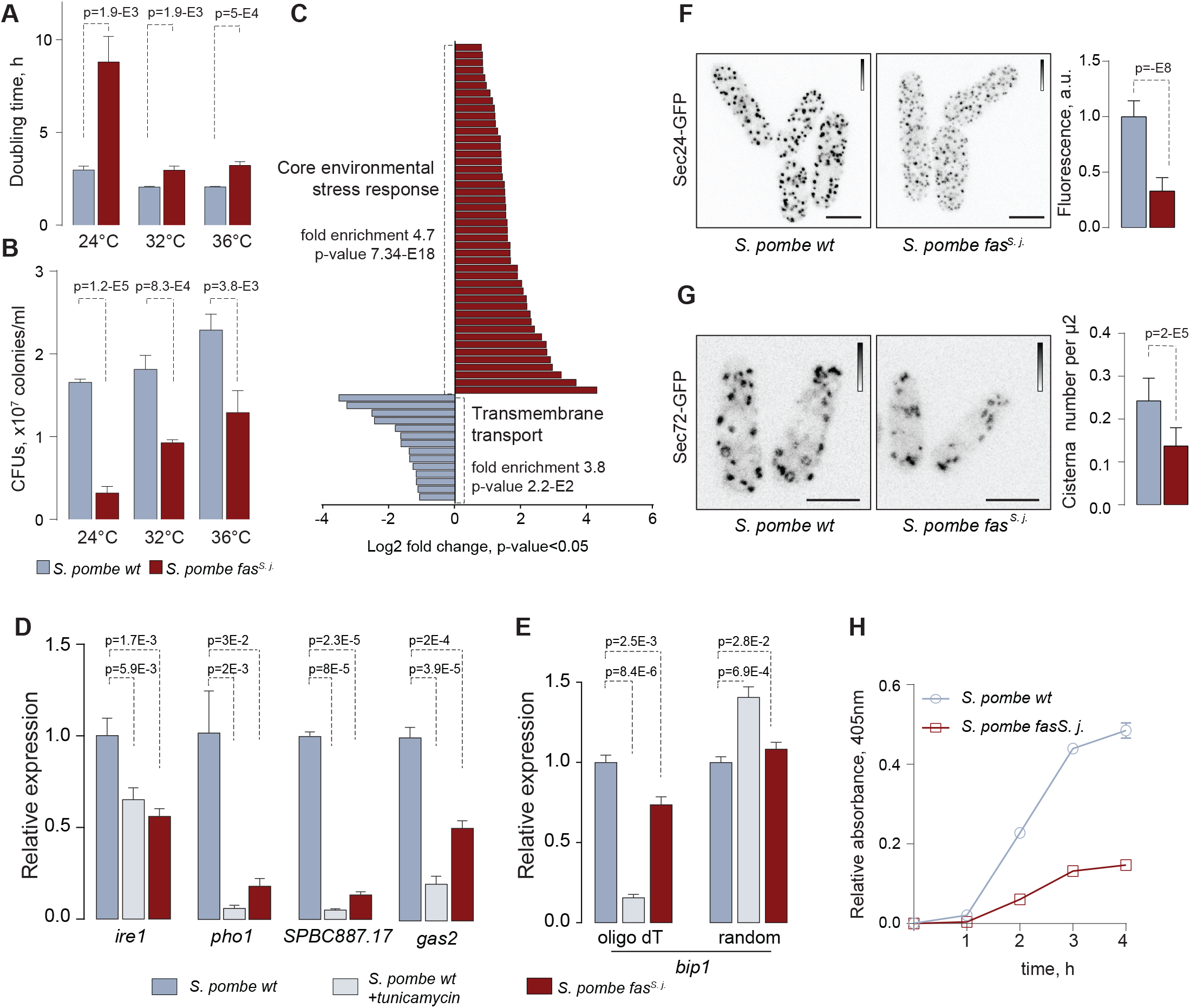
Genetic replacement of the cytosolic FAS complex in *S. pombe* with its S*. japonicus* counterpart results in UPR activation and defects in secretory pathway. (**A**) Doubling times for *S. pombe* wild type and *fas^s. j.^* cultures grown at indicated temperatures in the YES medium (*n = 3*). (**B**) A graph representing the survival of the *S. pombe* wild type and *fas^s. j.^* cells at indicated temperatures, in CFUs/ml of culture. (**C**) Differentially expressed genes constituting two largest functional categories in *S. pombe fas^s. j.^* cells as compared to the wild type. Cells were grown in YES at 24°C. AnGeLi suite was used for GO annotations. (**D**) A graph summarizing the results of qPCR analyses of indicated poly-A mRNAs in *S. pombe* wild type, *S. pombe* wild type treated with 0.5 µg/ml tunicamycin for one hour to induce the UPR and *S. pombe fas^s. j.^* mutant cells. (**E**) A graph showing the results of qPCR analyses of *bip1* poly-A and total mRNA species in *S. pombe* wild type, *S. pombe* wild type treated with 0.5 µg/ml tunicamycin for one hour and *S. pombe fas^s. j.^* mutant cells. (**D**-**E**) Shown are the mean values derived from three biological and two technical repeats, normalized to the wild type. *p* values derived from the unpaired parametric t-test. (**F**) Maximum intensity z-projections of spinning-disk confocal images of *S. pombe* wild type and *fas^s. j.^* cells expressing Sec24-GFP. Average fluorescence intensities relative to the wild type are presented on the right. (n = 142 and 154 cells for the wild type and the mutant, respectively). (**G**) Maximum intensity z-projections of spinning-disk confocal images of *S. pombe* wild type and *fas^s. j.^* cells expressing Sec72-GFP. For each genotype, trans-Golgi cisternae numbers per unit of area are presented on the right. (n = 153 and 167 cells for the wild type and the mutant, respectively). (**H**) A graph demonstrating the efficiency of acid phosphatase secretion measured in *S. pombe* wild type and *fas^s. j.^* cells. Shown are the mean values ± SD (*n* = 3). (**F**, **G**) Scale bar, 5 μm. Calibration bars are shown for each image.

*S. pombe fas^s. j.^* cells exhibited global changes in mRNA abundance as compared to the wild type, as assessed by RNAseq, both at 24°C and 30°C. Differentially expressed genes were broadly comparable between the two growth conditions although we also detected some potential temperature-specific regulation events (Supplemental Fig. 3E and Supplemental Tables 8-9). The largest group of upregulated transcripts included the core environmental stress response (CESR) genes (Fig. 3C; Supplemental Fig. 3F and Supplemental Tables 8-9). Yet another group of upregulated mRNAs was linked to FA metabolism, including malonyl-CoA production and FA elongation (29.1 fold enrichment, p=0.008; Supplemental Table 8).

Of note, we observed a striking enrichment of the gene ontology (GO) term ‘transmembrane transport’ among significantly downregulated genes (Fig. 3C, Supplemental Fig. 3F and Supplemental Tables 8-9). This suggested a possibility of a global membrane-related cellular response to the FAS substitution. Indeed, several downregulated genes, including *pho1* encoding a major secreted acid phosphatase [17], were previously identified as targets of the RIDD-type unfolded protein response (UPR) in *S. pombe*. In this organism, a transmembrane ER-resident kinase/endonuclease Ire1 governs the decay of ER-associated mRNAs, working together with the downstream mRNA no-go-decay machinery [18, 19]. An RT-qPCR-based expression analysis of a number of Ire1 targets, including those identified in our RNAseq datasets, revealed a significant decrease in their steady-state mRNA levels in *S. pombe fas^s. j.^* cells as compared to the wild type (Fig. 3D and Supplemental Fig. 3G).

As expected, similar, albeit more pronounced downregulation was observed in cells treated with the inducer of ER stress, tunicamycin (Fig. 3D and Supplemental Fig. 3G). Unlike other UPR targets, the *bip1* mRNA is stabilized by the cleavage of its poly-A tail, supporting activity of this major chaperone upon ER stress [18]. In line with this, we observed a decline in poly-A *bip1* mRNA but an overall message stabilization in *S. pombe fas^s. j.^* mutants (Fig. 3E). Importantly, *ire1* deletion was synthetically lethal with *fas^s. j.^* replacement. Together, these data are consistent with the hypothesis that substituting the *S. pombe* FAS with the *S. japonicus* complex leads to a chronic activation of the UPR and that survival of the retroengineered cells relies on the presence of this ER homeostasis pathway.

We reasoned that the presence of the unusual C10:0-containing asymmetrical membrane GPLs in *S. pombe fas^s. j.^* cells may impact on the organization and function of the secretory pathway, where lipid composition is tightly regulated [20]. Indeed, we observed drastic downregulation of transitional ER (tER) in mutant cells, as visualized by the tER marker Sec24-GFP [21] (Fig. 3F). Furthermore, the *fas^s. j.^* mutant cells exhibited a reduced number of trans-Golgi cisterna (Fig. 3G). Consistent with UPR-dependent downregulation of *pho1* mRNA (Fig. 3D) and general defects in ER and Golgi organization, *S. pombe fas^s. j.^* cells were severely defective in acid phosphatase secretion (Fig. 3H). Thus, introduction of asymmetrical phospholipids to *S. pombe* cells leads to pronounced defects in functions of the secretory pathway.

Membrane composition and physical properties may impose constraints on the structure of transmembrane parts of integral membrane proteins [22]. The comparisons of predicted transmembrane helices (TMHs) in all orthologous groups of transmembrane proteins between *S. pombe* and other fission yeast species (1:1:1:1 orthologs) showed that predicted *S. japonicus* TMHs contained many small non-polar amino acids and fewer large non-polar residues compared to the rest of fission yeasts (Supplemental Fig. 4A-C). Interestingly, the pairwise comparison of predicted TMHs in single-TMH proteins revealed a statistically significant enrichment of relatively short TMHs in *S. japonicus* (Fig. 4A and Supplemental Table 10). For a number of *S. japonicus* proteins, TMHs failed to be predicted with confidence altogether (Fig. 4B and Supplemental Table 10). Thus, it appears that transmembrane domains of at least some orthologous proteins in *S. japonicus* diverged greatly compared to those in other fission yeasts. Such differences might be a result of structural adaptation of TMHs in *S. japonicus* to profound changes in the FA composition of membrane phospholipids (Fig. 1).

**Fig. 4.**
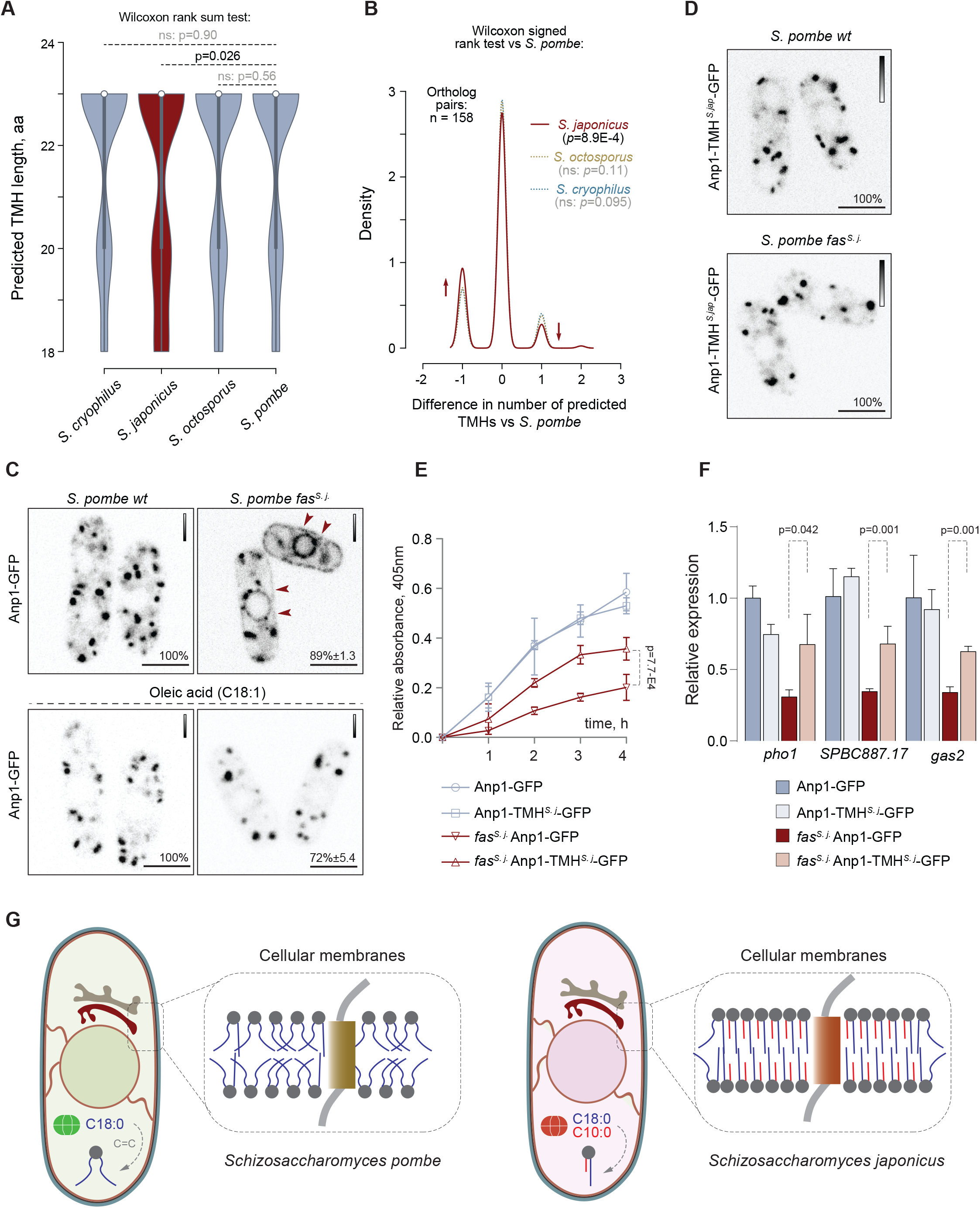
*Transmembrane* helices may have co-evolved with membrane lipids in *S. japonicus*. (**A**) A violin plot for distributions of predicted TMH lengths in the four fission yeast species. 1:1:1:1 orthologs of single-span TMH proteins were selected for the analysis. *p* values derived from the Wilcoxon rank sum test. (**B**) A density trace graph summarizing pairwise differences in the number of predicted TMHs in *S. japonicus*, *S. octosporus* and *S. cryophilus*, as compared to *S. pombe*. Only 1:1:1:1 orthologs of *S. pombe* single-span TMH proteins were included in the analysis. *p* values derived from the Wilcoxon rank sum test. (**C**) Single plane spinning-disk confocal images of cells expressing Anp1-GFP in *S. pombe* wild type and *fas^s. j.^* cells in the absence (n=205 and n=153 cells, respectively) or the presence of C18:1 supplementation (n=109 and n=97 cells, respectively). (**D**) Single plane spinning-disk confocal images of cells expressing Anp1-TMH^S.j.^-GFP as a sole copy of the protein (107 cells with endogenous FAS and 110 cells with replaced FAS were counted). (**E**) A graph summarizing the results of the acid phosphatase secretion assay in *S. pombe* cultures of indicated genotypes. Shown are the mean values ± SD (*n* = 3). (**F**) A graph summarizing the results of qPCR analyses of indicated poly-A mRNAs. Shown are the mean values derived from three biological and two technical repeats, normalized to the wild type. *p* values derived from the unpaired parametric t-test. (**G**) A diagram summarizing our hypothesis on co-evolution of transmembrane helices and membrane lipids in *S. pombe* and *S. japonicus*. (**C**, **D**) Scale bar, 5 μm. Calibration bars are shown for each image. Included are the percentages of cells in a population exhibiting the indicated phenotypes. (**E**, **F**) *p* values derived from the unpaired parametric t-test.

We reasoned that *S. pombe* proteins, orthologs of which exhibit TMH shortening or poor TMH prediction in *S. japonicus* as compared to other fission yeast species, might be particularly vulnerable to the introduction of the *S. japonicus* FAS to *S. pombe* and hence, an increase in GPL acyl chain asymmetry. To test this hypothesis, we visualized the subcellular localization of four TMH-divergent candidates, predicted to localize to various membrane sub-compartments. A mannosyltransferase complex subunit Anp1 that normally localizes to *cis*-Golgi [21], was mis-targeted in *S. pombe fas^s. j.^* cells, partitioning clearly between the ER and Golgi (Fig. 4C). Correction of cellular FA composition by exogenous oleic acid (C18:1) led to the reversal of this mis-localization pattern (Fig. 4C). We have observed a similar phenotype with the Anp1-interacting protein Gmh5/Mnn10 [23] (Supplemental Fig. 4D), suggesting that the localization and function of this entire complex could be affected. Curiously, the hydrophobic helix in *S. japonicus* Gmh5 does not reach the TMH prediction threshold, unlike its *S. pombe* counterpart (Supplemental Table 10). Similar mis-targeting phenotypes were observed for two other protein candidates that were predicted to have a shorter TMH or failed to reach the TMH prediction cutoffs in *S. japonicus*, including the ER-lipid droplet diacylglycerol transferase Lro1 [24] and the ER-plasma membrane tethering protein Tcb2 [25] (Supplemental Fig. 4D).

The predicted TMH in Anp1 was 18 amino acids in *S. japonicus* as compared to 20 amino acids in *S. pombe* and 23 amino acids in other fission yeasts (Supplemental Fig. 4E). Despite this difference, endogenous Anp1 proteins exhibited a typical Golgi localization in both *S. pombe* and *S. japonicus* (Fig. 4C and Supplemental Fig. 4F). To establish if mis-localization of Anp1 in *S. pombe fas^s. j.^* cells could be driven by a mismatch between its TMH structure and the rewired membrane lipid composition, we engineered an Anp1 chimerae in *S. pombe* by replacing the native TMH with its shorter *S. japonicus* version and keeping the rest of the protein intact (Supplemental Fig. 4E). Strikingly, this was sufficient to restore the Golgi localization pattern of Anp1 in *S. pombe fas^s. j.^* cells (Fig. 4D). The swap did not affect Anp1 Golgi localization in the wild type, indicating functional versatility of the *S. japonicus* TMH.

We did not observe significant differences in the growth rates between the Anp1-GFP and the Anp1-TMH*^S. j.-^*GFP *S. pombe fas^s. j.^* cultures (Supplemental Fig. 4G). Yet, recovering Anp1 Golgi localization by the TMH swap led to a significant improvement in acid phosphatase secretion efficiency in *S. pombe fas^s. j.^* cells (Fig. 4E). Since *pho1* is a target of the UPR pathway (Fig. 3D), we tested if this could be due to a higher abundance of the *pho1* mRNA. Indeed, the steady state levels of *pho1* poly(A) mRNA and other UPR targets were partially rescued in *S. pombe fas^s. j.^* cells expressing Anp1-TMH*^S. j.-^*GFP (Fig. 4F). Thus, restoring the localization of the Anp1-containing mannosyltransferase complex to Golgi is sufficient to ameliorate at least some of the defects induced by grafting the *S. japonicus* FAS to *S. pombe*.

We have shown that *S. japonicus*, a sister species to a well-studied model organism *S. pombe*, synthesizes abundant structurally asymmetrical phospholipids, where the medium acyl chain C10:0 is linked to the *sn*-2 position of the glycerol backbone and the *sn*-1 position is occupied by a long, usually saturated FA (Fig. 1). *In vitro*, membranes assembled from *S. japonicus* phospholipid mixtures are more ordered and consistently, are characterized by a higher bending rigidity (Fig. 2) [26]. At least two factors may contribute to such a difference. First, the FA tails of *S. japonicus* phospholipids are considerably more saturated (Fig. 1E) and therefore are likely to form closely packed and more ordered bilayers. Another possibility is that a large difference in the lengths of two acyl chains of individual phospholipid molecules may lead to partial interdigitation of FAs, leading to an increase in the strength of lateral interactions and tighter packing. In the gel phase, the 8-carbon acyl length asymmetry in pure PC assemblies does result in longer FAs spanning the entire bilayer and an overall decrease in membrane thickness [27, 28]. However, even if interdigitation occurs in liquid phase, it is likely extremely dynamic since the translational diffusion of lipids is not affected and the in-plane elasticity is increased in membranes made of asymmetrical phospholipids compared to their symmetrical equivalents [29, 30]. *In vivo*, biophysical properties of the membrane may be further modified by other components including sterols and sphingolipids. Co-existence of asymmetrical and symmetrical phospholipids in *S. japonicus* (Fig. 1) may potentially enhance the formation of functionally relevant membrane domains. Further work will be required to assess how the asymmetrical lipids are partitioned within the cell and the bilayer and how this may impact on cellular physiology.

The molecular changes underlying production of large amounts of C10:0 alongside the LCFAs and assembly of asymmetrical phospholipids might have allowed *S. japonicus* to colonize new ecological niches. One of the most important biophysical parameters, membrane fluidity, is usually adjusted to different growth temperatures by modulating the extent of cellular FA desaturation and sterol output [31, 32]. However, eukaryotic FA desaturation and sterol biosynthesis are oxygen-demanding processes, and this presents a problem in anaerobic conditions. Curiously, anaerobically-grown budding yeast replace a large proportion of their ‘normal’ phospholipids containing long FA tails with structurally asymmetrical molecules where the shorter C10:0, C14:0 and, in particular, C12:0 are confined to the *sn-*2 position of the glycerol backbone [33]. Similarly, defective desaturation of long FAs appears to trigger an increase in the synthesis of medium chain FAs even in aerobic conditions [34]. This suggests that phospholipid structural asymmetry may provide an alternative way of maintaining membrane physico-chemical properties when FA desaturation and/or sterol synthesis is not possible. In line with possible adaptation to anaerobic lifestyle, *S. japonicus* does not respire and grows well in the absence of oxygen, unlike *S. pombe* [35, 36]. It also undergoes robust transitions between yeast and hyphal forms, suggesting that in the wild this organism may have a burrowing, hypoxic lifestyle [37, 38]. Additional benefits of assembling tightly packed membranes from largely saturated lipids may include reduced permeability for oxygen and potentially hazardous solutes encountered in the environment [39–41].

Both polypeptide chains of fungal cytosolic FA synthases tend to exhibit overall sequence conservation (see [42] for an excellent review). The *S. japonicus* FAS is not unusual in this respect (see Experimental Procedures for the corrected *fas1* ORF), although it does have a number of potentially interesting substitutions that should be analyzed in the future. Of note, mutations in the ketoacyl synthase domain of the budding yeast *fas2* can shift the product spectrum from long to medium chain FAs [43]. The *S. japonicus* FAS is different in this respect as it exhibits two sharply defined product peaks rather than an overall shift towards shorter FAs. Yet, the budding yeast mutagenesis data suggest that the evolutionary acquisition of new functionality may not necessarily require a large number of steps.

Importantly, other cellular factors may influence the spectrum of FAS products. Although the *S. japonicus* FAS produces C10:0 *in vitro* and when expressed in *S. pombe*, the proportion of this medium chain FA in these circumstances are lower than in the native cellular context (Fig. 2F-G). Our data did not support the hypothesis that C10:0 production in *S. japonicus* might require the acyl-CoA-binding protein proposed to control FA termination (Supplemental Fig. 2L) and [16, 44]). An interesting alternative could be the different ratios between acetyl- and malonyl-CoA in *S. japonicus* and *S. pombe*. It is known that relative depletion of malonyl-CoA results in the enzyme favoring priming over elongation [42] and increases the production of shorter FAs. Curiously, the mRNA for Cut6, an enzyme responsible for malonyl-CoA synthesis [45], is significantly upregulated in *S. pombe* cells expressing the *S. japonicus* FAS (Supplemental Tables 8 and 9), perhaps as a part of an endogenous compensation mechanism triggered by elevated C10:0 levels. Other cellular adaptations are likely required to efficiently handle C10:0 alongside the long chain FAs, including modifications of lipid remodeling enzymes.

Replacement of the *S. pombe* FAS with its *S. japonicus* counterpart caused poor growth, especially at lower temperature and an activation of the core environmental stress response (Fig. 3A-C and Supplemental Fig. 3A, 3E-F). Such a signature may be attributed not only to downregulation of a large number of membrane transporters (Supplemental Tables 8-9) but also the changes in membrane biophysical properties, which could affect signaling through the cellular stress pathways [46, 47]. Importantly, FAS replacement in *S. pombe* induced chronic UPR (Fig. 3D-E and Supplemental Fig. 3G). The UPR in fission yeast operates through selective decay of mRNAs encoding proteins destined for the secretory pathway [18, 19]. We note that although the C10:0-containing molecules constitute less than 20% total phospholipids (Fig. 2G) this is sufficient to induce the UPR, suggesting that either the heterologous lipids are preferentially partitioned to the ER or that early secretory compartments are particularly vulnerable to lipid disbalance. Activation of Ire1 in *S. pombe fas^s. j.^* cells may be sustained by microdomain formation by the asymmetrical C10:0 containing lipids in the vicinity of Ire1 oligomers, which could lower the energy costs of membrane bending associated with oligomer formation [48]. Regardless of whether Ire1 senses bilayer stress directly [49–51], *S. pombe fas^s. j.^* cells depend on Ire1 for survival suggesting that UPR likely functions in this system to alleviate protein mis-folding and / or mis-targeting defects caused by changes in the FA composition (Fig. 3 and Supplemental Fig. 3). Given the reliance of *S. japonicus* on C10:0-containing largely saturated phospholipids, it will be interesting to see if this organism might have rearranged the structural features of Ire1 allowing it to sense ER lipid composition.

Our bioinformatics analyzes indicate that many *S. japonicus* transmembrane proteins might have experienced the need to adapt to changes in cellular lipid composition by reducing an average residue volume or shortening and/or remodeling the TM domains (Fig. 4A-B and Supplemental Fig. 4A-C). TMDs span the hydrophobic core of the lipid bilayer and therefore, specific membrane FA composition may impose constraints on their sequences. TMDs of single-spanning ER and *cis*-Golgi proteins tend to be shorter than those residing in the later compartments of the secretory pathway [22]. Curiously, most single-pass TM proteins that exhibit shortened TMDs in *S. japonicus* as compared to other fission yeasts are predicted to function largely at the ER, ER-organelle contacts and *cis*-Golgi (Supplemental Table 10). The membranes of the broad ER territory tend to exhibit looser lipid packing [11] and thus, their protein residents could be particularly sensitive to an increase in membrane order. In addition to reducing the hydrophobic length, other properties of the TMDs may facilitate their insertion and/or folding in well-ordered membranes – perhaps explaining the statistically significant increase in poorly predicted TMHs in *S. japonicus* single-pass TM proteins vs their 1:1:1:1 orthologs in other fission yeasts (Fig. 4B).

Supporting the hypothesis that TMHs co-evolve with membranes, the replacement of the TMH in *S. pombe* Anp1 with a shorter *S. japonicus* version rescues not only its mistargeting in *S. pombe* cells expressing the *S. japonicus* FAS (Fig. 4D), but also partially relieves downregulation of UPR targets in this genetic background (Fig. 4E-F). Anp1 is a key player in the assembly of the mannosyltransferase complex, which takes place in the ER before trafficking to Golgi [23, 52]. It is possible that ER retention of this complex may impact folding of proteins destined for secretion either through improper glycosylation events or contribute to ER stress indirectly. Yet, given the complexity of observed cellular phenotypes, the physiological consequences of introducing the *S. japonicus* FAS to *S. pombe* cells are likely not limited to this particular pathway.

Our work provides new insights into the generation of phospholipid diversity in evolution and its impact on cellular biology connected to membrane functions. We show that evolutionary changes in membrane FA composition may necessitate rapid adaptation of the transmembrane domains (Fig. 4G). The two fission yeast species with their divergent lifestyles constitute an ideal experiment of Nature, study of which may lead to a conceptually new understanding of the relationship between the underlying metabolic makeup and the evolution of cellular properties.

## Experimental Procedures

### Molecular genetics and strain husbandry

We used standard fission yeast media and methods [53–55]. Strains used in this study are listed in Supplemental Table 11. All primers are shown in Supplemental Table 12. Molecular genetics manipulations were performed using PCR- or plasmid-based homologous recombination using plasmids carrying *S. japonicus ura4*, kanR, hygR or natR cloned into the pJK210-backbone (pSO550, pSO820 and pSO893, respectively). Mitochondrial FA synthesis mutants *etr1Δ*, *mct1Δ* and *htd2Δ* of *S. japonicus* were generated using standard PCR-based recombination with KanMX6 as a selection marker. To generate a parental *S. pombe* strain carrying replacement of the two-component FAS complex with its *S. japonicus* counterpart we first replaced the *fas1* open reading frame (ORF) by homologous recombination, using a construct containing the 5’- and 3’-UTRs of f*as1^S.pombe^* that flanked the *fas1 ORF^S.japonicus^* and the kanMX6 cassette. Using similar strategy and the construct carrying the hygR cassette as a selection marker, we then replaced the *fas2* ORF. Of note, we have identified several mis-annotations in the *S. japonicus* gene encoding the FAS β subunit (SJAG_04107). Using both RNAseq / *de novo* transcript assembly and sequencing of cDNA obtained from vegetatively grown cells we identified the following errors: 1) T at position 1240 is absent; 2) an additional C is present at position 2158; 3) both annotated introns are in fact retained. The corrected sequence will be uploaded to EnsemblFungi. Replacements of the TMH in *S. pombe* Anp1 with its *S. japonicus* version were constructed similarly. All constructs were verified by Sanger sequencing. A PCR-based method [56] was used to tag Anp1, Gmh5, Lro1 and Tcb2 at the C-terminus, typically using KanMX6 or natR as a selection marker. Mating of *S. pombe* strains was performed on YPD solid medium and *S. japonicus*, on SPA solid medium. Spores were dissected and germinated on YES agar plates.

Detailed protocols of lipidomics, biochemical, biophysical, cell biological and physiological experiments are provided in Supplemental Experimental Procedures.

## Acknowledgments

We are grateful to the Oliferenko lab for discussions and D. Owen (King’s College London) for suggestions on the manuscript. Many thanks are due to V. Christodoulou (The Francis Crick Institute) for help with FAS purification, A. Le Marois (The Francis Crick Institute) for advice on GUV preparation and M. Grininger (University of Frankfurt) for advice on FAS assays. Our work was supported by the Wellcome Trust Senior Investigator Award (103741/Z/14/Z) to S. O.

## Author contributions

M. Makarova conceived and performed cell biological, biochemical and biophysical experiments, generated strains, analyzed data and co-wrote the manuscript. M. Peter, G. Balogh, A. Glatz and L. Vigh designed, performed and interpreted all lipidomics experiments except GC-MS, J. I. MacRae performed and analyzed the GC-MS experiments. N. Lopez Mora and P. Booth performed and interpreted measurements of membrane bending rigidity. E. Makeyev performed bioinformatics analyses. M. P., G. B., A. G., L. W., J. I. M., N. L-M. and E. M. edited the manuscript. S. Oliferenko conceived and interpreted experiments and wrote the manuscript.

## Data availability statement

The data that support the findings of this study are available from the corresponding author upon request. The authors declare that all data reported in this study are available within the paper and its supplementary information files.

## Supplemental information

Delineating the rules for structural adaptation of membrane-associated proteins to evolutionary changes in membrane lipidome

Maria Makarova, Maria Peter, Gabor Balogh, Attila Glatz, James I. MacRae, Nestor Lopez Mora, Paula Booth, Eugene Makeyev, Laszlo Vigh and Snezhana Oliferenko

## Supplemental Experimental Procedures

### Cell growth and survival measurements

For growth rate measurements, cells were grown in EMM (unless otherwise stated) overnight, diluted to OD_595_ 0.2 and supplemented with cerulenin (10µg/ml in DMSO) and/or FAs conjugated with BSA based on recommendations described in [1]. FA stock (5 mM) was prepared in 50% ethanol heated to 70°C. The FA solution was then conjugated with FA-free BSA at a 5:1 ratio at 37°C. Decanoic acid was used in methylated form. Final FA concentration added to the cell culture was 250µM, mixtures of FAs were added as separate stocks to the final concentration of 250µM. Control cells were treated with BSA solution. For survival measurements cells were grown in indicated conditions for 24 hours followed by serial dilutions and plating in triplicates on YES solid medium. The colony numbers were calculated after 48 hours of growth using OpenCFU software [2].

### ESI-MS-based lipidomic analysis

Lipid standards were from Avanti Polar Lipids (Alabaster, AL, USA). Solvents for extraction and MS analyses were liquid chromatographic grade (Merck, Darmstadt, Germany) and Optima LC-MS grade (Thermo Fisher Scientific, Waltham, MA, USA). All other chemicals were the best available grade purchased from Sigma-Aldrich (Steinheim, Germany). Phospholipase A2 (Snake venom from *Crotalus atrox*) was from Serva.

Exponentially growing yeast cell cultures in EMM were harvested and disrupted in water using a bullet blender homogenizer (Bullet Blender Gold, Next Advance, Inc., Averill Park, NY, USA) in the presence of zirconium oxide beads (0.5 mm) at speed 8 for 3 min at 4°C. Protein concentration of cell homogenates was determined using the Micro BCA™ Protein Assay Kit (Thermo Fisher Scientific, Waltham, MA, USA). A portion (40 µL of ∼500 µl total volume) of the homogenate was immediately subjected to a simple one-phase methanolic (MeOH) lipid extraction [3]. First, the homogenate was sonicated in 1 ml MeOH containing 0.001% butylated hydroxytoluene (as antioxidant) in a bath sonicator for 5 min, then shaken for 5 min and centrifuged at 10,000 x*g* for 5 min. The supernatant was transferred into a new Eppendorf tube and stored at −20 °C until MS measurement (∼25 nmol/ml total lipid in MeOH).

Electrospray ionization mass spectrometry (ESI-MS)-based lipidomic analyses were performed on a LTQ-Orbitrap Elite instrument (Thermo Fisher Scientific) equipped with a robotic nanoflow ion source TriVersa NanoMate (Advion BioSciences, Ithaca, NY) using chips with the diameter of 5.5-µm spraying nozzles. The ion source was controlled by Chipsoft 8.3.1 software. The ionization voltages were +1.3 kV and −1.9 kV in positive and negative mode, respectively, and the back-pressure was set at 1 psi in both modes. The temperature of the ion transfer capillary was 330°C. Acquisitions were performed at the mass range of 350-1300 *m/z* at the mass resolution Rm/z 400 = 240,000.

The lipid classes phosphatidylcholine (PC), lysophosphatidylcholine (LPC), diacylglycerol (DG), triacylglycerol (TG) and ergosteryl ester (EE) were detected and quantified using the positive ion mode, while phosphatidylethanolamine (PE), phosphatidylinositol (PI), phosphatidylserine (PS), their lyso derivatives LPE, LPI, LPS, phosphatidic acid (PA), phosphatidylglycerol (PG), cardiolipin (CL), ceramide (Cer), inositolphosphoceramide (IPC) and mannosyl-inositolphosphoceramide (MIPC) were detected and quantified using the negative ion mode.

For quantification, ∼20 µL lipid extract (corresponding to 1.2 µg protein) was diluted with 280 µL infusion solvent mixture (chloroform:methanol:isopropanol 1:2:1, by vol.) containing an internal standard mix (71 pmol PC d31-16:0/18:1, 7 pmol DG di22:1, 5 pmol TG tri22:1, 50 pmol PE d31-16:0/18:1, 43 pmol PI d31-16:0/18:1, 24 pmol PS d31-16:0/18:1, 1 pmol PG d31-16:0/18:1, 5 pmol PA d31-16:0/18:1, 1 pmol CL tetra14:0 (for *S. pombe*), 1 pmol CL tetra18:1 (for *S. japonicus*), 4 pmol Cer t18:0/16:0, and 18 pmol CE 18:1). Next, the mixture was halved, and 5% dimethylformamide (additive for the negative ion mode) or 3 mM ammonium chloride (additive for the positive ion mode) were added to the split sample halves. 10 µL solution was infused and data were acquired for 2 minutes.

MS/MS fragmentation experiments were carried out to determine FA compositions, ratio of isobaric species and FA positions. For fragmentation, HCD collision energy values were determined for PC, PE, PI and PS, and data-dependent MS/MS fragmentation experiments were performed based on mass lists from survey scans. To determine FA side chain regiochemistry, we followed the rule that losses of the neutral carboxylic acid ([M-H-RCOOH]−) or neutral ketene moieties ([M-H-RCH=C=O]−) are more abundant from the sn-2 position than from the *sn*-1 position [4].

Lipids were identified by LipidXplorer software [5] by matching the m/z values of their monoisotopic peaks to the corresponding elemental composition constraints. The mass tolerance was 3 ppm. Data files generated by LipidXplorer queries were further processed by custom Excel macros.

Lipid classes and species were annotated according to the classification systems for lipids [6]. For glycerolipids, the lipid class (*sn*-1/*sn*-2) format specifies the structures of the FA side chains as well as the side chain regiochemistry, e.g., PC (16:0/18:1). In sum formulas, e.g., PC (34:1), the total numbers of carbons followed by double bonds for all chains are indicated. For sphingolipids, the sum formula, e.g., Cer (44:0:4), specifies first the total number of carbons in the long chain base and FA moiety then the sum of double bonds in the long chain base and the FA moiety followed by the sum of hydroxyl groups in the long chain base and the FA moiety.

### Phospholipase A2 digestion

The total polar lipid fraction (TPL) was isolated from *S. japonicus* total lipid extract by thin-layer chromatography as described in [7]. Approximately 1 mg TPL isolate was dissolved in 250 µl diethyl ether, and then 250 µl borate buffer (0.1 M, pH 7.0), 125 µl CaCl2 (0.01 M) and 125 µl PLA2 solution (8 mg/ml PLA2 in water) were added; the mixture was stirred at room temperature in a test tube with glass stopper. After 2 h, 200 µl reaction mixture (corresponding to ∼100 µg TPL) was centrifuged at 2400 x*g*, the upper organic layer was evaporated, and the residue was extracted according to a modified Folch procedure [7]. The lower organic layer was evaporated under vacuum, and the residue was reconstituted in 1 ml chloroform:methanol 1:1 (v/v). The PLA2 digest was analysed together with the starting TPL isolate by ESI-MS as described above.

### DexPEG hydrogel films

DexPEG hydrogel films were prepared following previously reported methods [8]. Briefly, maleimide-modified dextran (1.5% weight solution) was cross-linked by PEG dithiol at room temperature. Typically for the preparation of 5 glass substrates with DexPEG hydrogel films, maleimide-modified dextran (75 mg) (degree of substitution = 3) was dissolved in water (4.5 g) and 23.6 mg of PEG dithiol (3400 Da) in water (0.5 g) were mixed to provide a hydrogel solution. The mixture was vortexed for 1 minute and 1 ml of hydrogel solution was immediately drop-casted on thiol functionalised microscope glass slides. The DexPEG substrates were stored at room temperature for further use.

### GUVs preparation in DexPEG hydrogel films

GUVs were prepared by the swelling of *S. pombe* or *S. japonicus* phospholipid mixtures on DexPEG hydrogel coated glass substrates at room temperature. Briefly, the lipids were dissolved in a mixture of methanol:chloroform (3:1) to a final concentration of 50 mg/ml. 20 µl of the desired lipid solution was then drop casted on DexPEG substrates and dried under a gentle stream of nitrogen gas for 5 minutes and placed under vacuum overnight. A growth chamber was made by placing a polydimethylsiloxane (PDMS) spacer between the hybrid lipid-DexPEG hydrogel coated slide and a microscope glass slide and clamped with crocodile clips. GUVs growth was initiated by hydrating the hybrid lipid-DexPEG film with a sucrose solution (400 µl, 100 mM sucrose). The hydrated substrates were left overnight at room temperature. Dense suspensions of GUVs were collected the following day and use for phase contrast microscopy.

### Phase contrast microscopy

GUVs suspensions (10 - 20 µl) were diluted in an observation chamber containing 400 µl of glucose solution (150 mM) for time-lapse microscopy imaging. GUVs were imaged in phase contrast mode with 1 ms exposure time and recorded for 40 seconds in a Nikon Eclipse TE-2000-E inverted microscope equipped with a digital high-speed camera Orca-Flash 4.0 (Hamamatsu) and a TI-DH Dia Pillar Illuminator (100W).

### Bending rigidity

The bending rigidity parameter was calculated by analyzing the thermal fluctuations of GUVs from the recorded time lapsed microscopy imaging with a home-made software and methodology as published previously [9, 10]. Fluctuation analyses were performed on 4,000 contours of individual GUVs. For small deformations, the distance from the membrane edge to the center of the vesicle about the mean edge position was Fourier-transformed to give a power spectrum that was adjusted between modes 6-20. The bending rigidity parameter was extracted from the fluctuation spectrum using Helfrich equation. A detailed mathematical description of the model can be found in [11, 12].

### Electroformation of GUVs and labelling with the FAST DIO

Chamber for GUV electroformation consisted of two conductive ITO glasses separated by a rubber spacer of 0.3 mm. Phospholipid extracts were dissolved in chloroform/methanol (2:1) and deposited on one of the ITO coated glasses. After drying the chamber was filled with 300 µl of 0.1 M Tris-HCl pH 7.5 and 200 mM sucrose. AC field of 10 Hz, 1 V was applied to the chamber. Electroformation was conducted at 40°C for 2 hours [13].

FAST DIO (Molecular Probes) was added at a concentration of 0.1 mol% to GUV mixtures followed by spinning disk confocal imaging at excitation 488 nm. Images were acquired with the same settings and intensities of the individual GUVs were analyzed at the equatorial plane.

### Metabolic labeling

For metabolic labeling experiments cells were grown in EMM overnight, diluted to OD_595_ 0.2 and supplemented with cerulenin (10 µg/ml in DMSO) and/or 200 µM stearic acid (C18:0; ^13^C labeled at each carbon atom) in 0.5% Brij35 for 4 hours. For glucose carbon atom tracing in newly synthesized FAs, glucose in the growth media was replaced with the uniformly labeled ^13^C glucose. Following 4-hour incubation, cells were collected and subjected to lipid extraction and sample drying.

### GC-MS

Dried samples, containing 10 nmol internal standard ([1-^13^C_1_]-lauric acid), were biphasic-partitioned using 700 µl chloroform:methanol:water (1:3:3, v/v). Data acquisition was performed using an Agilent 7890B-7000C GC-QQQ in EI mode after derivatization of dried apolar phase by the addition of 25 µl chloroform:methanol (2:1, v/v) and 5 µl MethPrep II (Grace). GC-MS parameters were as follows: carrier gas, helium; flow rate, 0.9 ml/min; column, DB-5MS (Agilent); inlet, 250°C; temperature gradient, 70°C (1 min), ramp to 230°C (15°C/min, 2 min hold), ramp to 325°C (25°C/min, 3 min hold). Scan range was m/z 50-550. Data was acquired using MassHunter software (version B.07.02.1938). Data analysis was performed using MANIC software, an in house-developed adaptation of the GAVIN package [14]. Fatty acid methyl esters (FAMES) were identified and quantified by comparison to authentic standards, and label incorporation estimated as the percentage of the metabolite pool containing one or more ^13^C atoms after correction for natural abundance.

### Fatty acid synthase purification

The procedure was adapted from [15], with some modifications. Cell were grown in YES media in a 40-liter fermenter for 24 hours up to OD_595_ 0.8, collected and washed with the buffer A (50 mM Tris-HCl pH 7.5, 200 mM NaCl, 1 mM EDTA, 1 mM TCEP), followed by resuspension in the same buffer with protease inhibitors (Pefabloc SC 1 mM, leupeptin 10 µg/ml, pepstatin A 10 µg/ml). Extracts were then frozen rapidly by dripping into liquid nitrogen. Freezer mill (SPEX CertiPrep 6850 Freezer/Mill) with 7 cycles of 2 min at crushing rate of 15 was used to disrupt cells and obtain powder for protein purification. Cell powder (15 g) was resuspended in buffer A with protease inhibitors to a final volume of 150 ml and placed on ice followed by ammonium sulfate protein precipitation. First, ammonium sulfate was added slowly with vigorous mixing at 0°C to the final concentration of 23%, followed by incubation for 30 minutes. Following centrifugation at 48000 x*g* for 40 minutes pellets were discarded. Ammonium sulfate was again added to the supernatant as described above, to the final concentration of 55%. Following centrifugation as above, protein pellets were resuspended in 200 ml of buffer A and subjected to anion exchange chromatography using the HiTrap Q HP 5 ml column. Protein fractions were collected from 100 to 800 mM NaCl and analyzed for the presence of FAS activity. Typically, fractions containing active FAS eluted within the 150-300 mM NaCl range. These active fractions were combined and concentrated to 1 ml using Amicon 10K ultra-filters. Protein concentrates were then loaded on a 10-40% linear sucrose density gradient in buffer A, prepared in 32S Beckman tubes. Those were centrifuged at 89000 x*g* for 18 hours at 4°C. Afterwards, 1 ml fractions were collected from top to bottom and assayed for FAS activity. Fractions containing concentrated active enzyme (typically, three) were yellow due to FAS binding to FMN [16]. To concentrate the activity and exchange buffer systems we then used MonoQ 5/50 GL anion exchange chromatography. 500 µl MonoQ fractions were collected and verified for FAS activity. Typically 1-2 fractions exhibited high activity and those were used for *in vitro* FA synthesis.

### FAS enzyme assay

Monitoring of NADPH oxidation by malonyl-CoA and acetyl-CoA at 340 nm was used to assess FAS activity for each step of FAS purification, as shown in [15]. The assay buffer consisted of 200 mM potassium phosphate pH 7.3, 50 µM acetyl-CoA, 25 µM malonyl-CoA, 100 µM NADPH, 2.5 mM EDTA, 4 mM dithiothreitol, and 0.03% bovine serum albumin. All components and the enzyme, except malonyl-CoA were mixed and a blank rate was measured for 20 minutes. The enzymatic reaction was then initiated by the addition of malonyl-CoA.

### Fatty acid synthesis in vitro

To analyze FAS product spectrum, active protein fractions after the MonoQ purification step were combined so that the total volume of the reaction was 400 µl. The buffered solution (200 mM potassium phosphate pH 7.3, 87.5 µM DTT, 2.25 mM NADPH, 0.20 mM acetyl-CoA, 1.00 mM malonyl-CoA) with added enzyme was kept at room temperature for 18 hours and the reactions were stopped by flash freezing.

### Fatty acid synthase assay – product analysis

FA-CoA assay products were extracted from the reaction mixture using a previously described protocol [15] with some modifications. Briefly, 4 volumes of ice-cold acetone and 3 nmol 13:0-CoA were added to 350 µl of FAS reaction mixture. The mixture was vortexed for 30 seconds, transferred for 1 hour to −20°C, and then centrifuged at 10,000 x*g* for 5 min at 4°C. The supernatant was evaporated under vacuum, and the residue was dissolved in 100 µl water by sonication for 5 min. The solution was centrifuged again at 10,000 x*g* for 5 min, the supernatant was transferred into a new Eppendorf tube, and stored at −20°C until mass-spectrometry analysis (which was performed within 2 days). For mass spectrometry measurements, the assay products were first diluted 50-fold with water, then a further 12-fold dilution was made by water:acetonitrile (containing 0.1% triethylamine) 1:1 (v/v). The diluted solutions were injected into the mass spectrometer in the negative ion mode at −1.5 kV ionization voltage at the mass resolution R_m/z 400_ = 240,000. For FA-CoA detection, the MS was programmed in SIM mode with 4 Da windows for the potential FA-CoAs, while MS/MS fragmentation experiments were performed at 25% normalized HCD energy. Quantitation and product profile determination was made based on the spiked 13:0-CoA amount. To exclude the possibility of non-enzymatic product formation, blank assay mixtures (i.e., FAS assay without malonyl-CoA) were analyzed in parallel. Fragments were identified based on [17].

### Image acquisition and analysis

Images of GUVs labelled with Fast DiO and live cells were obtained using Yokogawa CSU-X1 spinning disk confocal system mounted on the Eclipse Ti-E Inverted microscope with Nikon CFI Plan Apo Lambda 100X Oil N.A. = 1.45 oil objective, 600 series SS 488nm, SS 561nm lasers and Andor iXon Ultra U3-888-BV monochrome EMCCD camera controlled by Andor IQ3. Maximum intensity projections of 8–10 z-stacks of 0.5 µm step size images are shown in Fig. 3F-G and Supplemental Fig. 4F. For Fig. 4C-D and Supplemental Fig. 4D single plane images are presented. Fluorescence images are shown with inverted LUT (look-up table).

Image processing and quantifications were performed in Fiji [18]. Images within each experiment images are shown within the same display range. Calibration bars are included. For Fig. 3F average fluorescence intensity was measured per cell and normalized to the control. To measure the number of Golgi cisternae (Fig. 3G), maximum intensity projections of spinning disk confocal images were thresholded. Cisternae numbers were then normalized to cell area.

### RNA isolation

*S. pombe* and *S. pombe fas ^S. j.^* were grown overnight at 24°C or 30°C to OD_595_ 0.4. 10 OD units of cells were collected and subjected to total RNA extraction using a Qiagen RNeasy Plus Mini Kit.

### RNA sequencing

Strand-specific mRNA-seq libraries for the Illumina platform were generated and sequenced at the Advanced Sequencing Facility at the Francis Crick Institute. mRNA libraries were prepared using polyA KAPA mRNA Hyper_Prep with the subsequent quality control using Agilent Technology TapeStation system. A 50-cycle single-read sequence run was performed on the HiSeq 2500 Illumina instrument. Raw sequence data was mapped to the *S. pombe* genome using a Galaxy server pipeline (https://usegalaxy.org/). Quality control was performed using the FastQ Groomer module, reads were mapped to the fission yeast genome using the Tophat module and differential gene expression was assessed by the Cuffdiff module. Functional enrichment analyses were performed using AnGeLi [19]. Genes with fold change log2 values higher than 0.8 or lower than –0.8 and p-values < 0.05 were considered for the subsequent analyses.

### Reverse Transcription and Real-Time Quantitative PCR (RT-qPCR)

Reverse transcription was performed at 50°C for 1 hour in a total reaction volume of 20 µl containing 2 µg RNA, 0.5 µg oligo (dT) or hexamer primers with a Transcriptor First Strand cDNA Synthesis Kit (Roche). qPCRBIO Probe Blue Mix Lo-ROX was used for the real-time qPCR assay with the primers generated using Primer3 software [20]. The real-time qPCR was performed on an Applied BioSystem in three biological and two technical replicates. qRT-PCR signal was normalized to GAPDH mRNA expression levels.

### Measurements of oxygen consumption

Cells were grown in EMM to early stationary phase (OD595=1). Typically, 60 ml of cell culture were centrifuged gently, washed once with fresh EMM and resuspended in 80 ml of EMM. The measurements were made using an HI98193 oximeter equipped with Hl764073 probe (Hanna Instruments). Readings were recorded for 15 minutes.

### Acid phosphatase secretion assay

Cell were grown in EMM until mid-log phase, washed with EMM and diluted to OD_595_ 0.1 in fresh medium for further growth at 30°C. Initial samples were measured at the time of resuspension and used as a reference for further measurements which were taken hourly. Typically, 200 µl of conditioned media was mixed with 200 µl of substrate solution (10mM p-nitrophenyl phosphate, 0.1 M sodium acetate, pH 4) and incubated at 30°C for 30 minutes. Reactions were stopped by addition of 200 µl of 1 M sodium hydroxide and absorbance measured at 405 nm [21].

### Bioinformatics

Fission yeast protein sequences showing 1:1:1:1 relationship in hierarchical orthologous groups were downloaded from the OMA browser (https://omabrowser.org/oma/home/) [22]. Transmembrane helices (TMHs) were predicted by TMHMM 2.0 (http://www.cbs.dtu.dk/services/TMHMM/) [23] using default parameters. Predictions differing from the median TMH length (23 aa) by ≥50% (i.e. ≥12 aa) were discarded. TMH amino acid composition was analyzed using ProtParam (https://web.expasy.org/protparam/) [24]. Sequences were aligned using the accurate (L-INS-i) option of MAFFT (https://mafft.cbrc.jp/alignment/software/) [25]. Data were analyzed and plotted in R [26].

### Quantification and statistical analyses

Lipidomics data were obtained from at least 5 replicates (biological and technical) and *p*-values were derived using unpaired parametric heteroscedastic t-tests. Experiments shown in Fig. 2 were performed in three biological replicates and *p*-values were derived using unpaired parametric t-test. For Fig. 3, unpaired nonparametric Kolmogorov-Smirnov tests was used to compare bending rigidities and phase separation in GUVs. For qPCR data p-values were calculated using unpaired parametric t-test. Statistical analyses were performed in Prism 7 (GraphPad Software) and R [26].

## Supplemental Figures and Legends

**Supplemental Fig. 1, related to Fig. 1.**
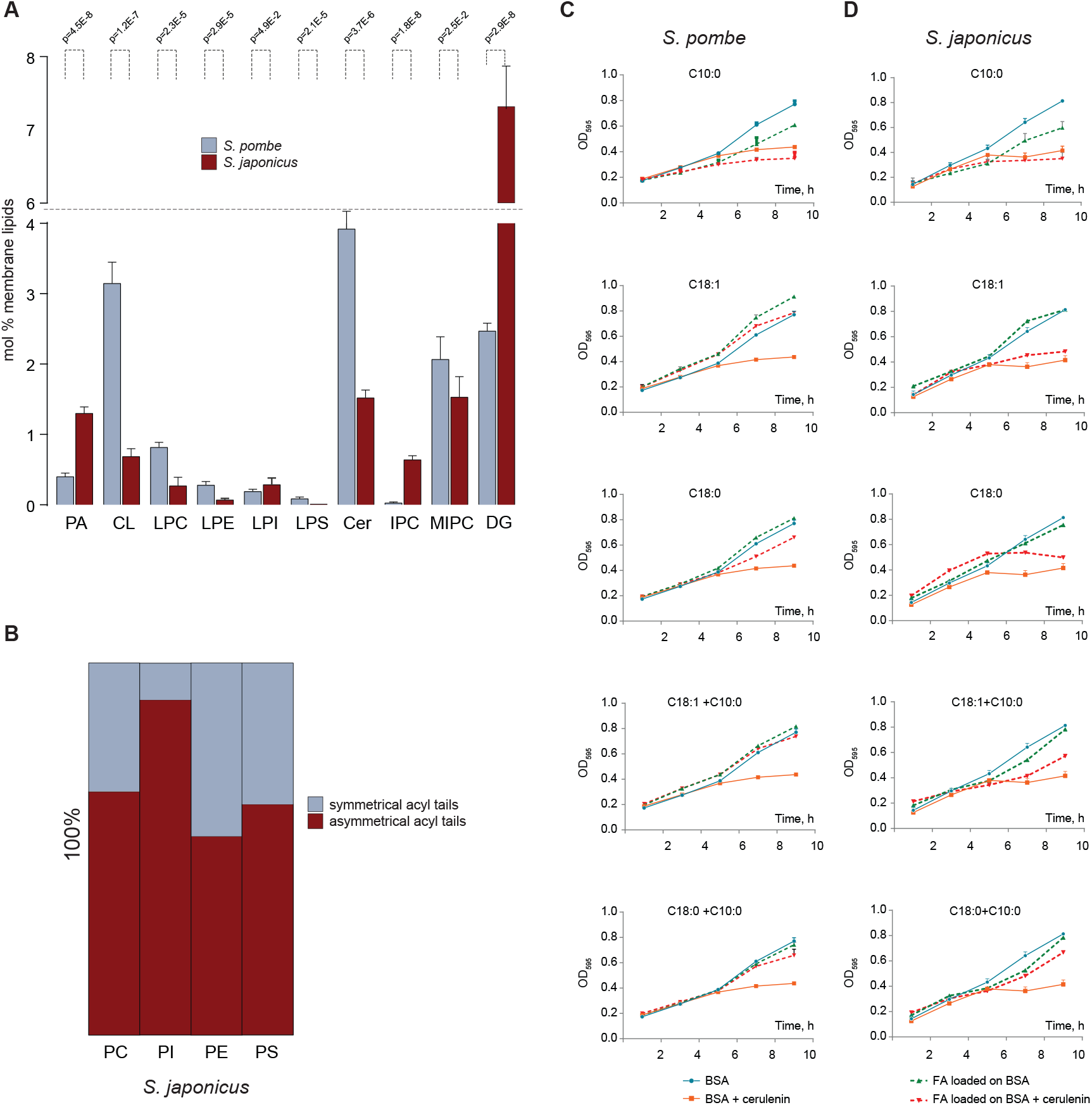
(**A**) Relative abundance of the indicated lipid classes in *S. pombe* and *S. japonicus*. Shown are the mean values ± SD (*n* = 5). *p* values derived from the unpaired parametric t-test. (**B**) A graph representing percentages of ‘asymmetric’ molecular species for the four indicated lipid classes in *S. japonicus*, calculated from the mass-spectrometry fragmentation analysis. Growth curves for *S. pombe* (**C**) and *S. japonicus* (**D**) cultures used to derive growth rates presented in Fig. 1I. (**C**-**D**) Shown are the mean values ± SD per time point (*n* = 3).

**Supplemental Fig. 2, related to Fig. 2.**
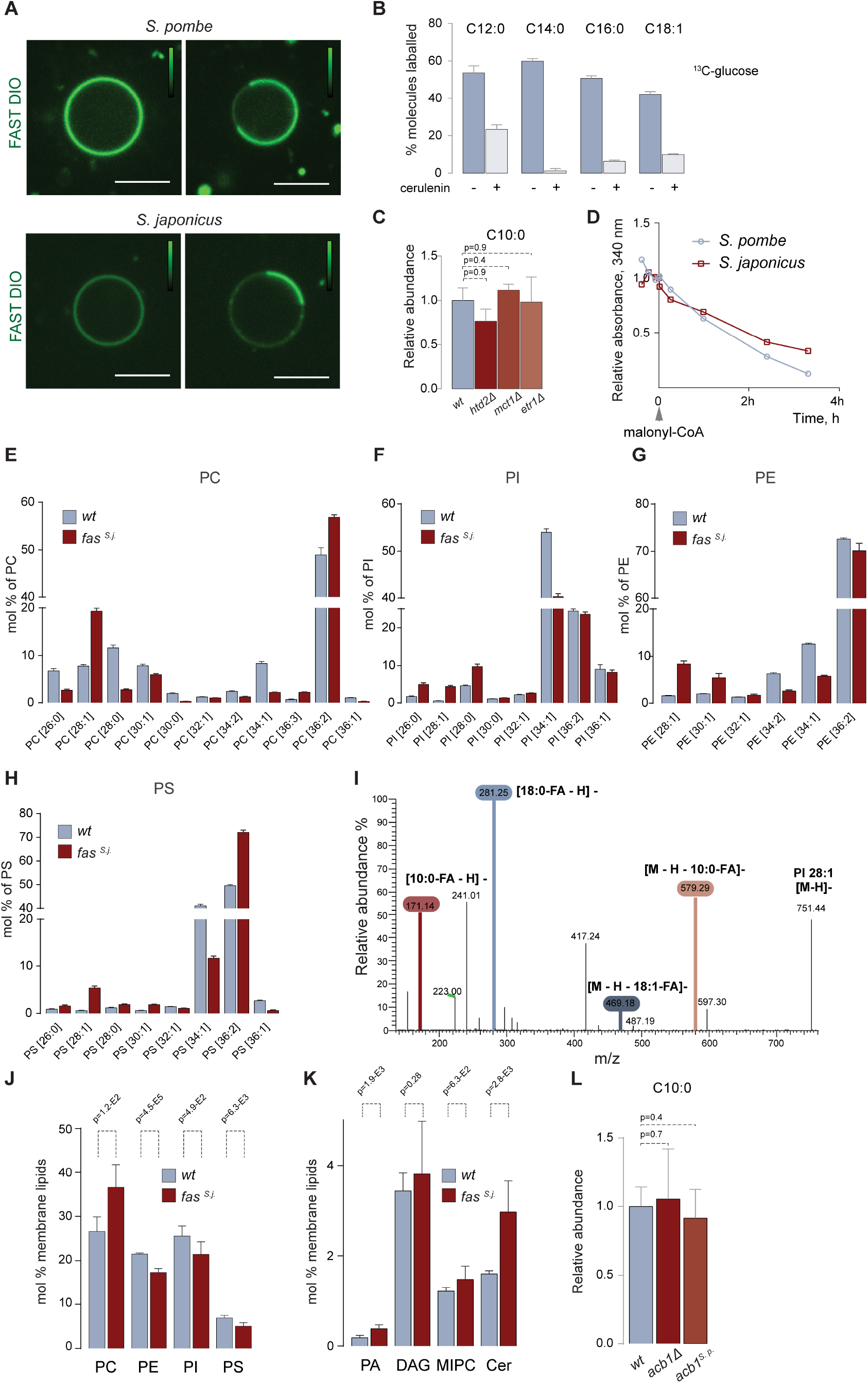
(**A**) Examples of the primary microscopy data used to calculate Ld/Lo values shown in Fig. 2B. Note differences in FAST DIO staining intensities between GUVs formed from *S. pombe* and *S. japonicus* phospholipids – this dye preferentially partitions to Ld membranes. Intensity calibration bars are included. Scale bar, 5 μm. (**B**) A plot describing the results of the GC-MS analysis of indicated ^13^C-labeled FAs extracted from *S. japonicus* cells grown in the presence of the U-^13^C-glucose for 4 hours. 10 μM cerulenin was added when indicated. Shown are the means of percentages of ^13^C-labeled fatty acids ± SD (n = 6). (**C**) Relative abundance of C10:0 in the wild type and three mutants in the mitochondrial FA synthesis pathway. Genotypes of mutants are indicated. (**D**) A line chart tracing relative absorbance of NADPH at 340 nm in protein fractions used for *in vitro* FA synthesis reactions shown in Fig. 2F. Malonyl-CoA was added at time 0. Graphs representing molecular species profiles for phosphatidylcholine (PC) (**E**), phosphatidylinositol (PI) (**F**), phosphatidylethanolamine (PE) (**G**) and phosphatidylserine (PS) (**H**) in *S. pombe* wild type and *S. pombe fas^s. j.^* cells. Phospholipids are specified by the total carbon atoms: total double bonds in acyls. Broken columns are used to show details at the bottom and the top of the scale. (**I**) A representative mass spectrum of the PI 28:1 molecular species from the *S. pombe fas^s. j.^* lipid extract indicating ions generated upon fragmentation. Graphs representing the abundance of the four main GPL classes (PC, PI, PE, and PS) (**J**) and other indicated membrane lipids (**K**) in *S. pombe* wild type and *fas^s. j.^* cells, presented as molecular percentages of membrane lipids. (**L**) Relative abundance of C10:0 in the wild type and *acb1* mutants of indicated genotypes. (**C**, **L**) Shown are the means of relative C10:0 content ± SD (n = 6). *p* values derived from the unpaired parametric t-test. (**E**-**H, J-K**) Shown are the mean values ± SD (*n* = 4 for the wild type and 5 for *fas^s. j.^* cells). *p* values derived from the unpaired parametric t-test.

**Supplemental Fig. 3, related to Fig. 3.**
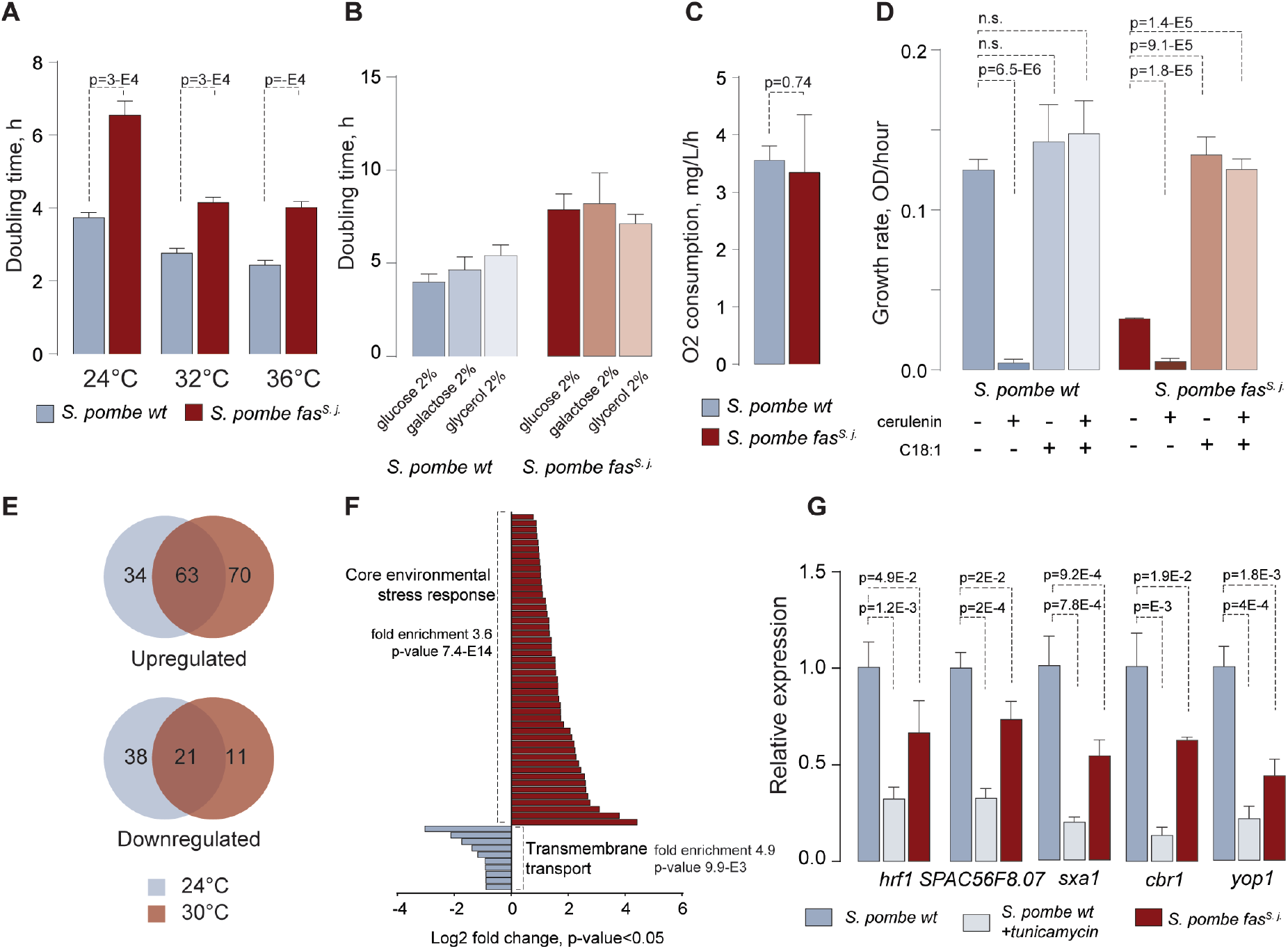
(**A**) Doubling times for *S. pombe* wild type and *fas^s. j.^* cultures grown at indicated temperatures in the EMM medium. (**B**) Doubling times for *S. pombe* wild type and *fas^s. j.^* cultures grown at 32°C in the YE-based media with indicated carbon sources (*n = 3*). (**C**) Oxygen consumption of *S. pombe* wild type and *fas^s. j.^* cultures grown in EMM at 32°C. (**D**) Growth rates of *S. pombe* wild type and *fas^s. j.^* cultures in the presence or the absence of cerulenin and C18:1 FA, as indicated. (**A**, **C**, **D**). Shown are the mean values ± SD (n = 3). *p* values derived from the unpaired parametric t-test. (**E**) Venn diagrams showing an overlap between the differentially expressed genes in wild type and *fas^s. j.^* S*. pombe* cells, at 24°C and 30°C. (**F**) Differentially expressed genes constituting two largest functional categories in *S. pombe fas^s. j.^* cells as compared to the wild type. Cells were grown in YES at 30°C. AnGeLi suite was used for gene category annotations. (**G**) A graph summarizing the results of qPCR analyses of indicated poly-A mRNAs in *S. pombe* wild type, *S. pombe* wild type treated with 0.5 µg/ml tunicamycin for one hour to induce the UPR and *S. pombe fas^s. j.^* mutant cells. Shown are the mean values derived from three biological and two technical repeats, normalized to the wild type. *p* values derived from the unpaired parametric t-test.

**Supplemental Fig. 4, related to Fig. 4.**
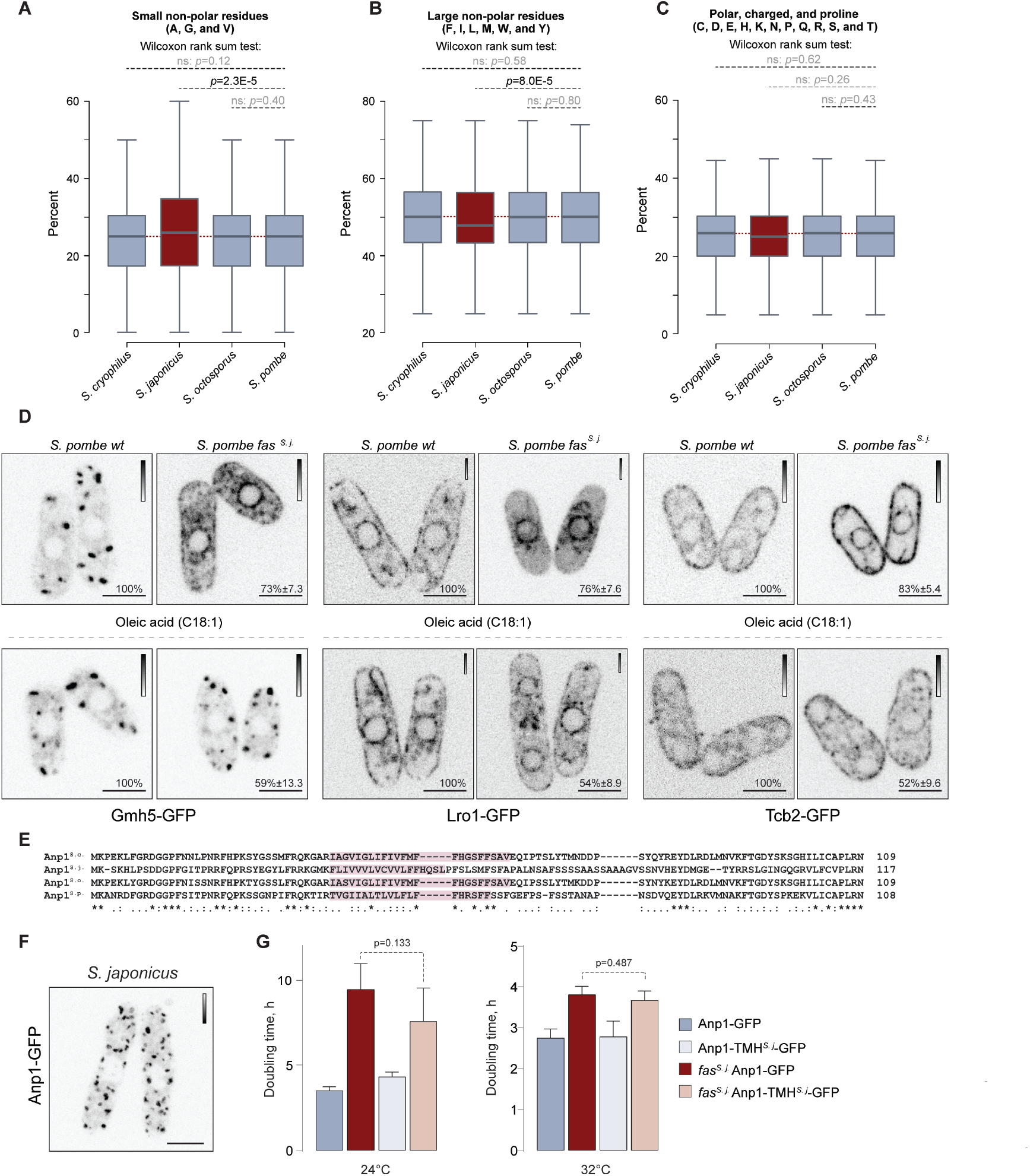
Box plots showing distributions of percentages of small non-polar residues (A), large non-polar residues (B) and the rest of amino acids (C) in all predicted TMHs in proteomes of the four fission yeasts. The dashed lines mark positions of the medians in *S. pombe*. Outliers are not shown. *p* values derived from the Wilcoxon rank sum test. (D) Single plane spinning-disk confocal images of cells expressing Gmh5, Lro1-GFP and Tcb2-GFP in *S. pombe* wild type and *fas^s. j.^* cells in the absence or the presence of C18:1 supplementation. Included are the percentages of cells in a population exhibiting the indicated phenotypes. (E) Multiple sequence alignments for N-terminal portions of Anp1 orthologs from the four fission yeast species including Anp1*^S.c.^* (OMA ID SCHCR05123), Anp1*^S.j.^* (SCHJY01063), Anp1*^S.o.^* (SCHOY00891) and Anp1*^S.p.^* (SCHPO02701). “*” corresponds to invariant, “:” to strongly conserved, and “.” to weakly conserved residues, according to the Clustal convention. TMHMM-predicted transmembrane helices are highlighted in pink. (F) Multiple projection of the spinning-disk confocal z-stack of *S. japonicus* cells expressing Anp1-GFP. (G) Doubling times of *S. pombe* cultures of indicated genotypes grown at 24°C and 32°C in the YES medium. Shown are the mean values ± SD (*n = 3*). *p* values derived from the unpaired parametric t-test. (D, F) Scale bar, 5 μm. Calibration bars are shown for each image.

